# Alterations in the Mammary Gland and Tumor Microenvironment of Formerly Obese Mice

**DOI:** 10.1101/2023.06.14.545000

**Authors:** Genevra Kuziel, Brittney N. Moore, Grace P. Haugstad, Yue Xiong, Abbey E. Williams, Lisa M. Arendt

## Abstract

Obesity is a risk factor for breast cancer, and women with obesity that develop breast cancer have a worsened prognosis. Within the mammary gland, obesity causes chronic, macrophage-driven inflammation and adipose tissue fibrosis. To examine the impact of weight loss on the mammary microenvironment, mice were fed high-fat diet to induce obesity, then switched to a low-fat diet. In formerly obese mice, we observed reduced numbers of crown-like structures and fibrocytes in mammary glands, while collagen deposition was not resolved with weight loss. Following transplant of TC2 tumor cells into the mammary glands of lean, obese, and formerly obese mice, diminished collagen deposition and cancer-associated fibroblasts were observed in tumors from formerly obese mice compared to obese mice. When TC2 tumor cells were mixed with CD11b^+^CD34^+^ myeloid progenitor cells, collagen deposition within the tumors was significantly greater compared to when tumor cells were mixed with CD11b^+^CD34^-^ monocytes, suggesting that fibrocytes contribute to early collagen deposition in mammary tumors of obese mice. Overall, these studies show that weight loss resolved some of the microenvironmental conditions within the mammary gland that may contribute to tumor progression.

## INTRODUCTION

Global obesity rates are continuing to rise [1, 2]. Obesity significantly increases the risk for the development of hormone receptor positive breast cancer in postmenopausal individuals [3-5]. Breast cancer patients with obesity have a significantly worse prognosis and overall survival regardless of menopausal status or tumor subtype [6]. Breast tissue is a depot of subcutaneous adipose tissue, and a hallmark of obesity is the recruitment of macrophages to form crown-like structures (CLS) to remove lipid and necrotic adipocytes [7]. In obesity, mammary adipose tissue is also associated with increased collagen deposition and stiffness surrounding adipocytes [8, 9], and smooth muscle actin (SMA)^+^ myofibroblasts also have been observed [8]. The inflammatory and fibrotic microenvironment within obese mammary tissue may promote tumor growth. Further, breast tumors from patients with obesity demonstrated higher levels of desmoplasia, which is characterized by increased SMA^+^ cancer-associated fibroblasts (CAF) and collagen deposition, than breast tumors from lean patients [8], suggesting that obesity also impacts the breast tumor microenvironment.

Weight loss ameliorates multiple health conditions associated with obesity. Epidemiological studies have shown that weight loss may decrease the risk for breast cancer in women with obesity [10, 11]. In patients treated with bariatric surgery for weight loss, macrophage populations appear to switch from an inflammatory to an alternatively activated phenotype in subcutaneous white adipose tissue [12, 13], which may enhance tissue repair [14]. The impact of weight loss on adipose tissue fibrosis is less clear [15, 16]. Changes in inflammation and adipose tissue fibrosis following weight loss have not been investigated in the mammary gland. Further, limited mouse models have examined how weight loss affects tumor growth and the tumor microenvironment. The effects of weight loss before tumor growth on the mammary tumor microenvironment have yet to be examined.

Fibrocytes, which originate in the myeloid progenitor cell population of the bone marrow, have attributes of both macrophages and myofibroblasts and have been identified in diseases characterized by inflammation and fibrosis [17, 18]. Fibrocytes are identified in tissues using combinations of markers including CD34, CD11b, CD45, SMA, and collagen I [19, 20]. In mammary tissue from obese mice, we observed increased numbers of CD34^+^CD11b^+^ cells with the ability to form colonies that expressed myofibroblast markers *in vitro*, consistent with fibrocytes [21]. In a tumor model of the inflammation associated with obesity, we also identified elevated numbers of fibrocytes in developing mammary tumors [22]. In human breast tissue, CD34^+^ cells have been detected in the extracellular matrix surrounding breast lobules and low grade DCIS but are no longer detected when SMA^+^ myofibroblasts were increased surrounding high grade DCIS and invasive ductal carcinoma [23, 24], which is suggestive of differentiating fibrocytes. While recent single cell RNA sequencing studies have identified fibrocytes in lung tumors [25, 26], the role of fibrocytes in altering the tumor microenvironment in obesity has not been examined.

Here, we investigate how weight loss impacts mammary gland inflammation and collagen fibrosis, as well as tumor growth and development of the microenvironment, using a diet-induced obesity mouse model. We observed that weight loss resolves CLS and reduces fibrocytes within the mammary gland, but total macrophages numbers and collagen deposition are not different. Tumors that develop in the mammary glands of formerly obese mice grow at an intermediate rate to tumors from obese or lean mice but have a tumor microenvironment more similar to tumors from lean mice. Interestingly, fibrocytes were decreased in tumors of obese mice. However, transplant of estrogen receptor alpha (ERα)^+^ TC2 tumors mixed with myeloid progenitor cells from obese mice leads to lasting increases in collagen and CAF within tumors. Together, these results suggest that weight loss prior to tumor formation reduces desmoplasia within tumors, potentially through reduced numbers of fibrocytes prior to tumor progression.

## MATERIALS AND METHODS

### Transgenic Mice

All animal procedures were conducted in compliance with a protocol approved by the University of Wisconsin Institutional Animal Care and Use Committee and housed in AAALAC accredited facilities (Animal Welfare Assurance Number: D16-00239). FVB.Cg-Tg(CAG-EGFP)B5Nagy/J mice (EGFP; 003516) [27] were purchased from Jackson Laboratory (Bar Harbor, ME, USA). FVB/NTac female mice were purchased from Taconic Biosciences. Mice were given food and water *ad libitum* and weighed weekly. Three-week-old female mice were fed either a low-fat diet (LFD; 16% kcal from fat; 2920X; Teklad Global; ENVIGO) or a high-fat diet (HFD; 60% kcal from fat; Test Diet, 58Y1) for 16 weeks. After 16 weeks on the HFD, mice were switched to the LFD for 6 weeks to induce weight loss.

### Isolation of Cells

Bone marrow was flushed from the humerus and femurs. Mammary glands and tumors were mechanically minced then enzymatically dissociated for 1 hr (mammary gland) or 1.5 hr (tumor) at 37°C in DMEM (10-017-CV; Corning Inc., Corning, NY, USA) supplemented with 10% FBS, 1% antibiotic/antimycotic solution (30-004-CI; Corning, Inc.), 1.5 mg/mL collagenase A (11088793001; MilliporeSigma, Burlington, MA, USA), and 125 U/mL hyaluronidase (H3506; Sigma-Aldrich, St. Louis, MO, USA). Both bone marrow and mammary gland were treated with ACK Lysing Buffer (10-548E; Lonza, Basel, Switzerland) to lyse red blood cells. Mammary glands and tumors were dissociated to single cells as described [28].

### Cell Lines

Parental and GFP^+^ TC2 cells were obtained from the lab of Dr. Linda Schuler [29]. Parental TC2 cells were cultured in DMEM (10-017-CV; Corning Inc.) supplemented with 10% fetal bovine serum and 1% antibiotic/antimycotic solution (30-004-CI; Corning, Inc). GFP^+^ TC2 cells were cultured in complete media supplemented with 1 mg/ml Geneticin (G418 Sulfate) (11811023; ThermoFisher, Waltham, MA, USA). Cells were cultured at 37°C with 5% CO_2_.

### Flow Cytometry and FACS Isolation

Both bone marrow and mammary gland cells were prepared as previously described [22], with antibodies in Table S1. Flow cytometry was performed using a BD LSRFortessa (BD Biosciences; San Jose, CA, USA). Fluorescence-activated cell sorting (FACS) was performed using a BD FACS Aria III cell sorter (BD Biosciences; San Jose, CA, USA) at the Flow Cytometry Laboratory (Carbone Cancer Center, University of Wisconsin-Madison). Gates were set using fluorescence-minus-one (FMO) controls. Data were analyzed using FlowJo 10.8.1 (Becton, Dickinson and Company, Ashland, OR, USA).

### TC2 Mammary Tumor Transplant

50,000 GFP^+^ TC2 tumor cells were mixed with 1:1 DMEM : Matrigel (354234; Corning, Inc.) and injected into bilateral inguinal mammary glands of lean, obese, and formerly obese mice. For transplant of TC2 tumor cells mixed with bone marrow cells, FACS was used to isolate total CD45^+^ cells, CD45^+^CD11b^+^CD34^-^ monocytes, and CD45^+^CD11b^+^CD34^+^ myeloid progenitor cells from bone marrow of EGFP mice. 25,000 FACS-isolated bone marrow cells were mixed with 50,000 parental TC2 cells and injected into mammary glands of lean and obese mice. Tumor length and width were recorded once a week with calipers. Tumor volume was calculated using the formula (L*W*W)/2. Once tumors reached 1 cm in length, mice were humanely euthanized, and tumors were collected.

### Immunomagnetic Cell Sorting and Fibrocyte Culture

Immunomagnetic bead sorting was performed as previously described [22], with the following modifications. CD11b^+^ cells were plated at 20,000 cells/well on 12-well plates, in four experiments. Colonies were counted 5 days after plating. Fibrocyte colonies grown for immunofluorescent staining were plated on 8-well chamber slides (15434; ThermoFisher) coated with poly-L-lysine (PLL; P4707; Sigma-Aldrich) and grown for 10 days. All sorted cells were grown in Mouse MesenCult Expansion Kit Media (05513; StemCell Technologies, Vancouver, BC, Canada). Cells were cultured at 37 °C with 5% CO_2_. To image colonies, cells were fixed with cold 100% methanol for 20 minutes at -20 °C and then stained with 0.1% crystal violet. Colony images were captured using a Keyence BZ-X710 microscope (Itasca, IL).

### Histology and Immunofluorescence

Tissue was paraffin-embedded and sectioned by the Experimental Animal Pathology Laboratory (Carbone Cancer Center, University of Wisconsin-Madison). Picrosirius red staining was completed as described [30]. Immunohistochemistry and immunofluorescence were performed as described [31]. All antibodies are listed in Table S1. Tissue was snap frozen in O.C.T. compound (23-730-571; ThermoFisher) and sectioned. Frozen tissue sections were fixed in cold 100% methanol, then washed in PBS. Slides stained for quantification were blinded, then imaged with identical image acquisition settings using a Leica TCS SP8 Confocal Microscope (Leica Microsystems, Buffalo Grove, IL, USA) or a Keyence BZ-X710 microscope. Picrosirius red staining around mammary ducts and tumors was imaged and quantified as described [31]. Picrosirius red and SMA stained tumor sections were tile-scanned and stitched using BZ-X Analyzer Software 1.3.0.3 (Keyence). SMA staining was imaged with DAPI and quantified using ImageJ. A ratio was calculated for SMA area : total area. F4/80 staining within tumors was quantified using the Color Deconvolution 2 ImageJ plugin [32]. At minimum 5 images per mammary gland or tumors were quantified and the values were averaged.

### Statistical Analysis

Significance was determined at *p*-values of 0.05 or less. Data were tested for normality using the Shapiro-Wilk test prior to further statistical analysis. One-way ANOVA with Tukey’s multiple comparisons test was used unless stated. GraphPad Outlier Calculator was used to identify significant outliers. Error bars represent mean ± S.E.M. unless stated. Statistical analyses were conducted using GraphPad Prism 9.4.1 (GraphPad Software, San Diego, CA, USA).

## RESULTS

### Weight loss partially resolves inflammation but not fibrosis

We have previously observed that mice switched from HFD to LFD lose a significant amount of weight and resolve obesity-induced changes in mammary epithelial cell populations [28]. To investigate the effects of weight loss on stromal cells in the mammary gland, 3-week-old female FVB/N mice were fed either LFD or HFD for 16 weeks. A group of mice fed HFD was then switched to the LFD for 6 weeks to induce weight loss (Figure 1A). These mice rapidly lost a significant amount of weight (p<0.0008; Figure 1B). Within the mammary glands of formerly obese mice, we observed significantly reduced formation of CLS formed by both F4/80^+^ macrophages (p<0.0001; Figure 1C) and CD11b^+^ myeloid lineage cells (p<0.0001; Figure 1D) compared to HFD-fed mice. These results demonstrate a loss of this signature of inflammation with weight loss.

**Figure 1.**
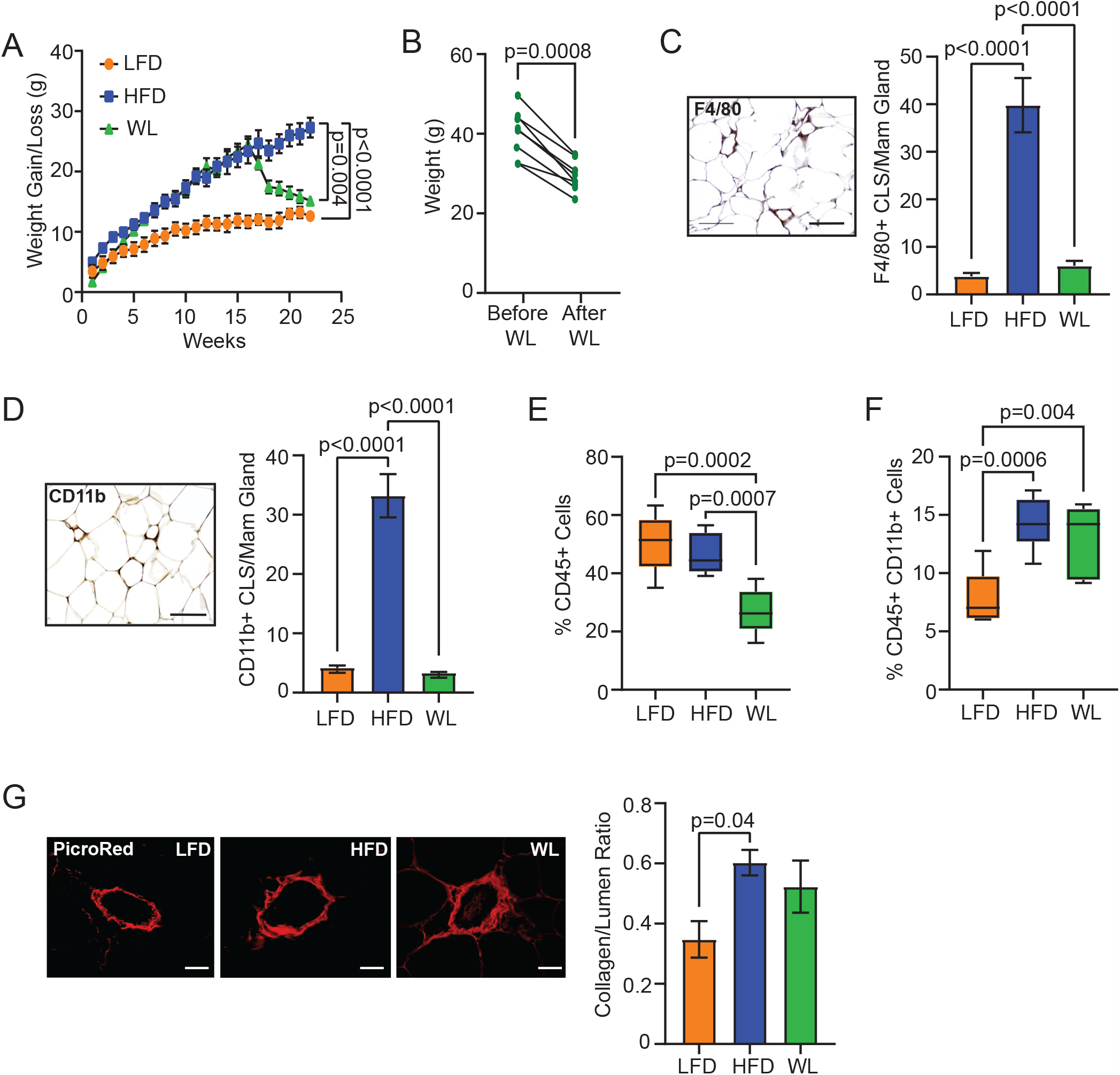
Weight loss partially resolves characteristics of obesity in mammary glands. **(A)** 3-week-old female FVB/N mice were fed LFD or HFD for 16 weeks, then a group was switched from HFD to LFD for 6 weeks (n = 8-10/group; Two-way ANOVA). **(B)** Weight change of mice switched from HFD to LFD for 6 weeks (n = 8 mice; paired t-test). **(C)** F4/80^+^ CLS per mammary gland section (n = 5-6/group). **(D)** CD11b^+^ CLS per mammary gland section (n = 5-6/group). **(E)** CD45^+^ cells in mammary glands of lean, obese, and formerly obese mice quantified by flow cytometry (n = 6-7/group). **(F)** CD45^+^CD11b^+^ myeloid cells in mammary glands of each group quantified by flow cytometry (n = 6-7/group). **(G)** Picrosirius red-stained collagen surrounding mammary ducts normalized to lumen size (n = 5 mice/group). Magnification bar: **(C, D)** 50 μm; **(G)** 25 μm.

To further examine how weight loss impacts myeloid lineage cells, mammary glands from lean, obese, and formerly obese mice were dissociated to single cells and stained with antibodies to detect CD45, CD11b, and CD34 by flow cytometry (Figure S1). Total CD45^+^ cells were decreased in formerly obese mice compared to either lean (p=0.0002) or obese mice (p=0.007; Figure 1E). However, total CD45^+^CD11b^+^ myeloid cells were significantly increased in both obese (p=0.0006) and formerly obese mice (p=0.004) compared to lean mice (Figure 1F). These data indicate that obesity induces an influx of myeloid cells into the mammary gland, which is not resolved after 6 weeks of weight loss.

Obesity increases fibrosis in the mammary gland [33, 34]. To determine if fibrosis is reduced with weight loss, collagen surrounding ducts was quantified using picrosirius red staining in mammary glands of lean, obese, and formerly obese mice. Collagen surrounding mammary ducts was significantly increased in obese mice compared to lean mice (p=0.04, Figure 1G), while collagen in formerly obese mice was at an intermediate level between obese and lean mice (Figure 1G). These results show that while weight loss resolved CLS, fibrosis was a longer-lasting part of the microenvironment.

### Weight loss decreases myeloid progenitor cells and fibrocytes

Fibrocytes have been associated with fibrosis in multiple contexts and are thought to originate in the myeloid progenitor cell population of the bone marrow [17]. To assess the effects of weight loss on myeloid progenitor cells, we quantified bone marrow cells from lean, obese and formerly obese mice using flow cytometry (Figure S2). CD45^+^CD11b^+^CD34^+^ myeloid progenitor cells were increased in bone marrow of obese mice compared to lean mice (p=0.009, Figure 2A), and weight loss reduced myeloid progenitor cells in formerly obese mice compared to obese mice (p=0.007, Figure 2A). Within the mammary gland, CD45^+^CD11b^+^CD34^+^ immature myeloid cells were significantly increased in obese mice (p=0.0002) and reduced in formerly obese mice compared to obese mice (p=0.002, Figure 2B).

**Figure 2.**
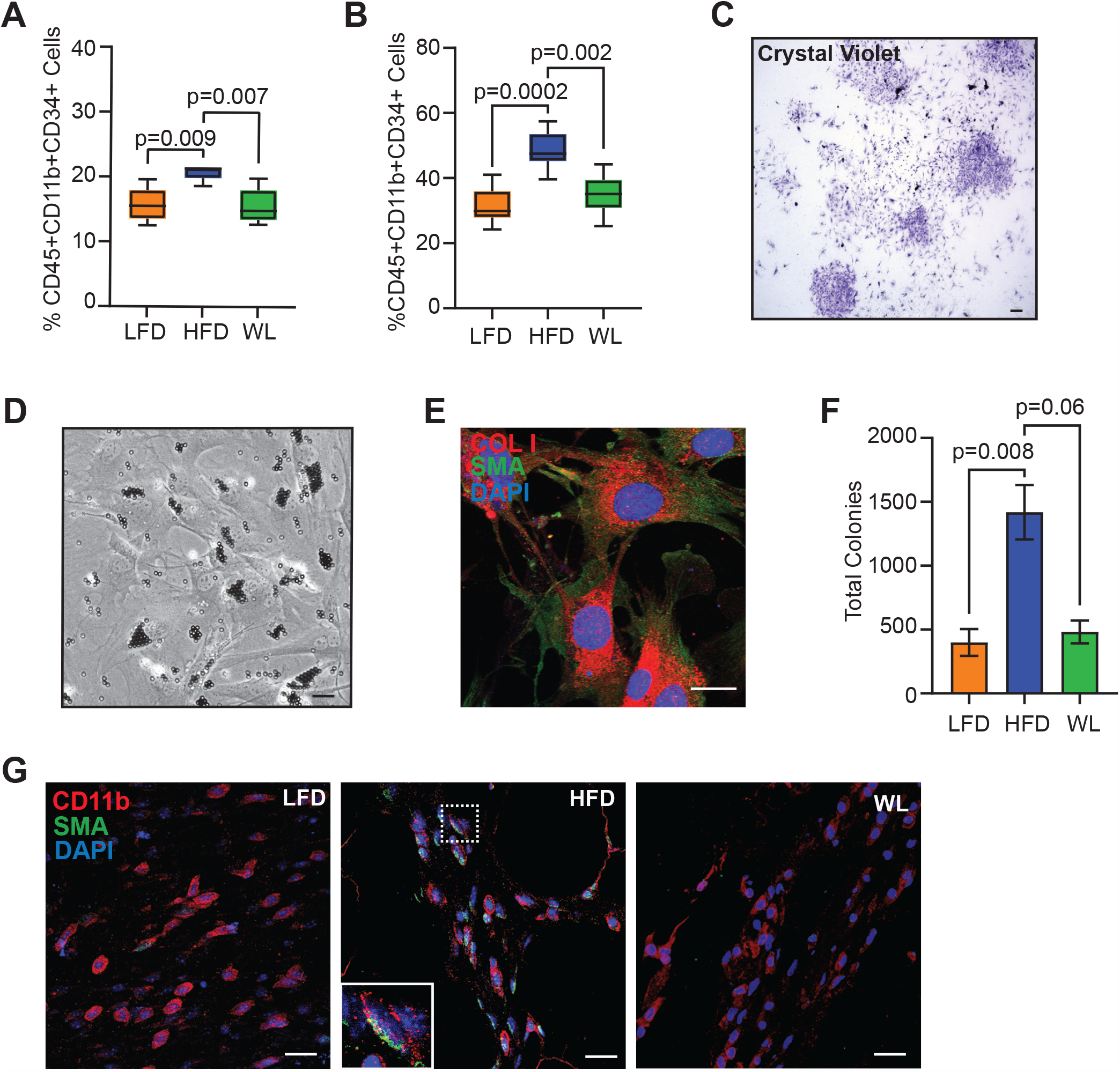
Weight loss reduces fibrocyte colony formation within the mammary glands. **(A)** CD45^+^CD11b^+^CD34^+^ myeloid progenitor cells in bone marrow quantified by flow cytometry (n = 5-6/group). **(B)** CD45^+^CD11b^+^CD34^+^ immature myeloid cells in mammary glands quantified by flow cytometry (n = 6-7/group). **(C)** Representative crystal violet image of colonies grown from immunomagnetically sorted CD11b^+^ cells. **(D)** Representative phase contrast image of an adherent colony grown from CD11b^+^ cells. **(E)** Representative image of a colony grown from CD11b^+^ cells stained to detect SMA and collagen I. **(F)** Colony quantification of CD11b^+^ sorted cells from mammary glands (n = 4/group). **(G)** Representative images of mammary glands stained for CD11b and SMA. Magnification bar: **(C, D)** 100 μm; **(E, F)** 25 μm.

We have previously shown that fibrocyte colonies form from CD11b^+^CD34^+^ cells [22]. To examine the impact of weight loss on fibrocytes, CD11b^+^ cells were immunomagnetically sorted from the mammary glands of lean, obese, and formerly obese mice, and assessed for their ability to form adherent colonies *in vitro* (Figure 2C). Fibrocytes formed adherent colonies that had a morphology similar to myofibroblasts (Figure 2D) and expressed markers SMA and collagen I (Figure 2E). CD11b^+^ cells isolated from obese mice generated significantly more fibrocyte colonies than lean mice (p=0.008), while fibrocyte colonies were reduced in CD11b^+^ cells from formerly obese mice compared to those from obese mice (p=0.06, Figure 2F).

To examine fibrocytes within tissue, mammary glands from lean, obese, and formerly obese mice were stained for CD11b and SMA (Figure 2G). Double positive cells were observed more frequently in the mammary glands of obese mice compared to either lean or formerly obese mice. Altogether, these results suggest that weight loss reduces fibrocyte numbers within mammary glands.

### Weight loss prior to tumor growth reduces tumor fibrosis

We have previously observed that obesity increases mammary tumor growth [35, 36]. To investigate the impact of weight loss on mammary tumor growth and the tumor microenvironment, we generated lean, obese, and formerly obese mice. Formerly obese mice weighed significantly less than obese mice (p=0.004), but significantly more than lean mice (p=0.04, Figure 3A). ERα^+^ TC2 mammary tumor cells were injected into the mammary glands of mice in each group. Mammary tumors grew significantly faster in obese mice compared to lean mice (p=0.0002, Figure 3B). Mammary tumors in formerly obese mice demonstrated an intermediate growth rate, but still grew significantly faster than tumors in lean mice (p=0.04, Figure 3B). Mammary tumors retained ERα expression at end stage (Figure 3C), and we have previously observed that growth in the mammary glands of obese mice did not change ERα expression compared to lean mice [36].

**Figure 3.**
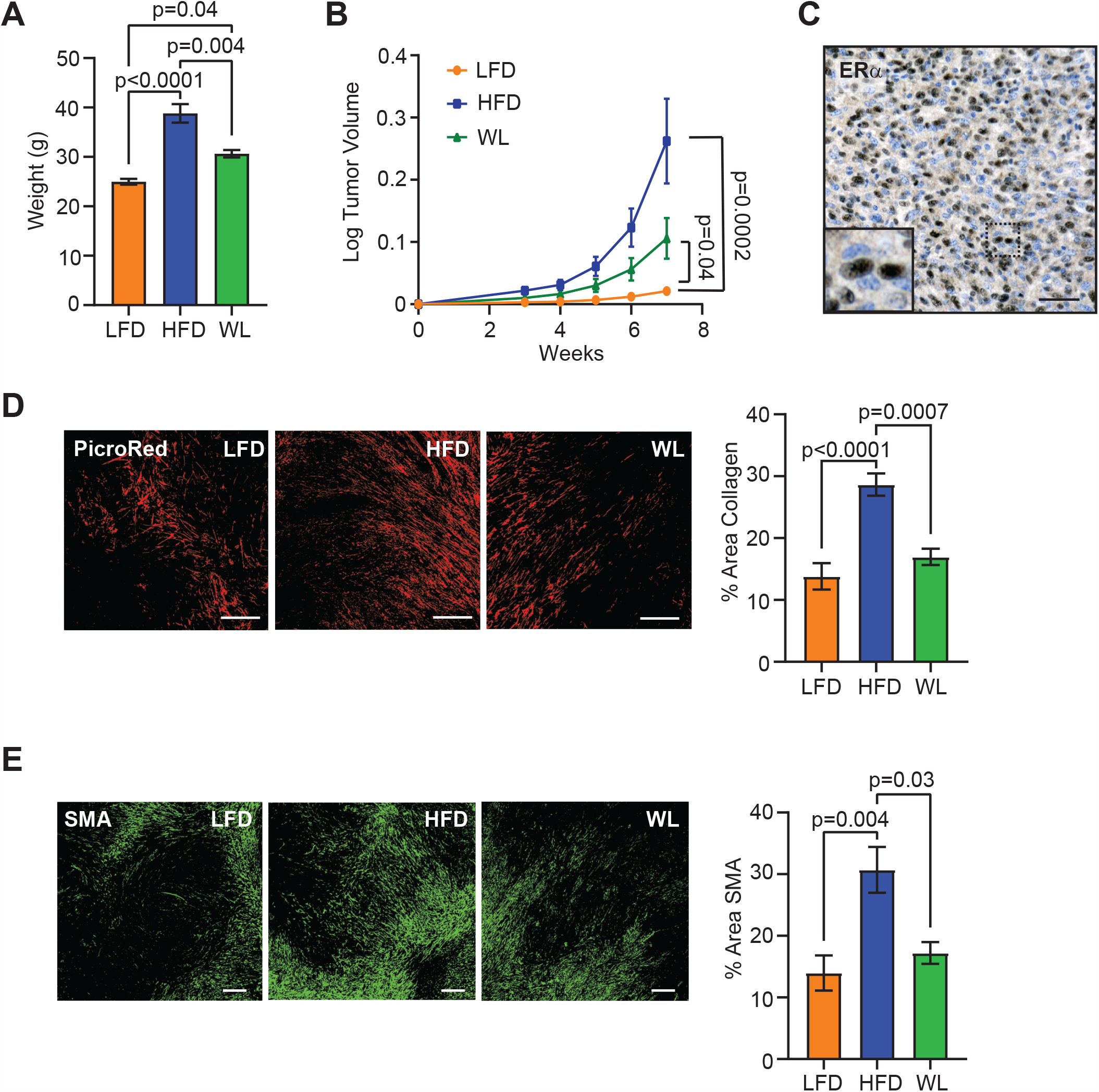
Weight loss preceding TC2 tumor cell transplant reduces CAF and collagen within tumor microenvironment. **(A)** Mouse weights at time of surgery (n = 8-10/group). **(B)** TC2 tumor growth following injection of LFD, HFD, and WL inguinal mammary glands (n = 8-10/group; Kruskal-Wallis with Dunn’s multiple comparisons test). **(C)** Representative image of ERα expression in TC2 mammary tumors. **(D)** Picrosirius red-stained collagen within tumors normalized to total tumor area (n = 6/group). **(E)** SMA expression within tumors normalized to total tumor area (n = 6/group). Magnification bar: **(C)** 50 μm; **(D, E)** 300 μm.

To assess how weight loss prior to tumor growth impacted the mammary tumor microenvironment, tumor sections from lean, obese, and formerly obese mice were stained with picrosirius red and collagen deposition was quantified. Collagen was significantly increased within tumors of obese mice compared to lean mice (p<0.0001, Figure 3D), and collagen within tumors of formerly obese mice was significantly diminished compared to obese mice (p=0.0007, Figure 3D). Tumor sections were also stained with SMA to identify CAF. Obesity significantly increased CAF in tumors compared to lean mice (p=0.004, Figure 3E). CAF were significantly reduced in tumors from formerly obese mice compared to obese mice (p=0.03, Figure 3E). Taken together, these data indicate that the mammary microenvironment after weight loss may no longer be primed to promote to collagen deposition and CAF formation in mammary tumors but still contributes to tumor growth.

### Weight loss prior to tumor growth reduces an immunosuppressive microenvironment

Since we observed that obesity and weight loss impacted the myeloid progenitor cell population in the bone marrow and immature myeloid cells in mammary gland of non-tumor-bearing mice (Figure 2A, B), we assessed how tumor growth altered these cells. We isolated bone marrow from tumor-bearing lean, obese, and formerly obese mice and analyzed the cells using flow cytometry. Similar to non-tumor-bearing mice, obese mice had significantly increased myeloid progenitor cells compared to lean mice (p=0.004, Figure 4A), and weight loss reduced myeloid progenitor cells compared to obese mice (p=0.02, Figure 4A). In contrast to the mammary glands of non-tumor-bearing mice, no differences were observed in the percentage of total CD45^+^CD11b^+^ myeloid cells in mammary tumors from mice in each group (Figure 4B). Further, CD11b^+^CD34^+^ immature myeloid cells were decreased in tumors of both obese mice (p=0.02) and weight loss mice (p=0.04) compared to lean mice (Figure 4C).

**Figure 4.**
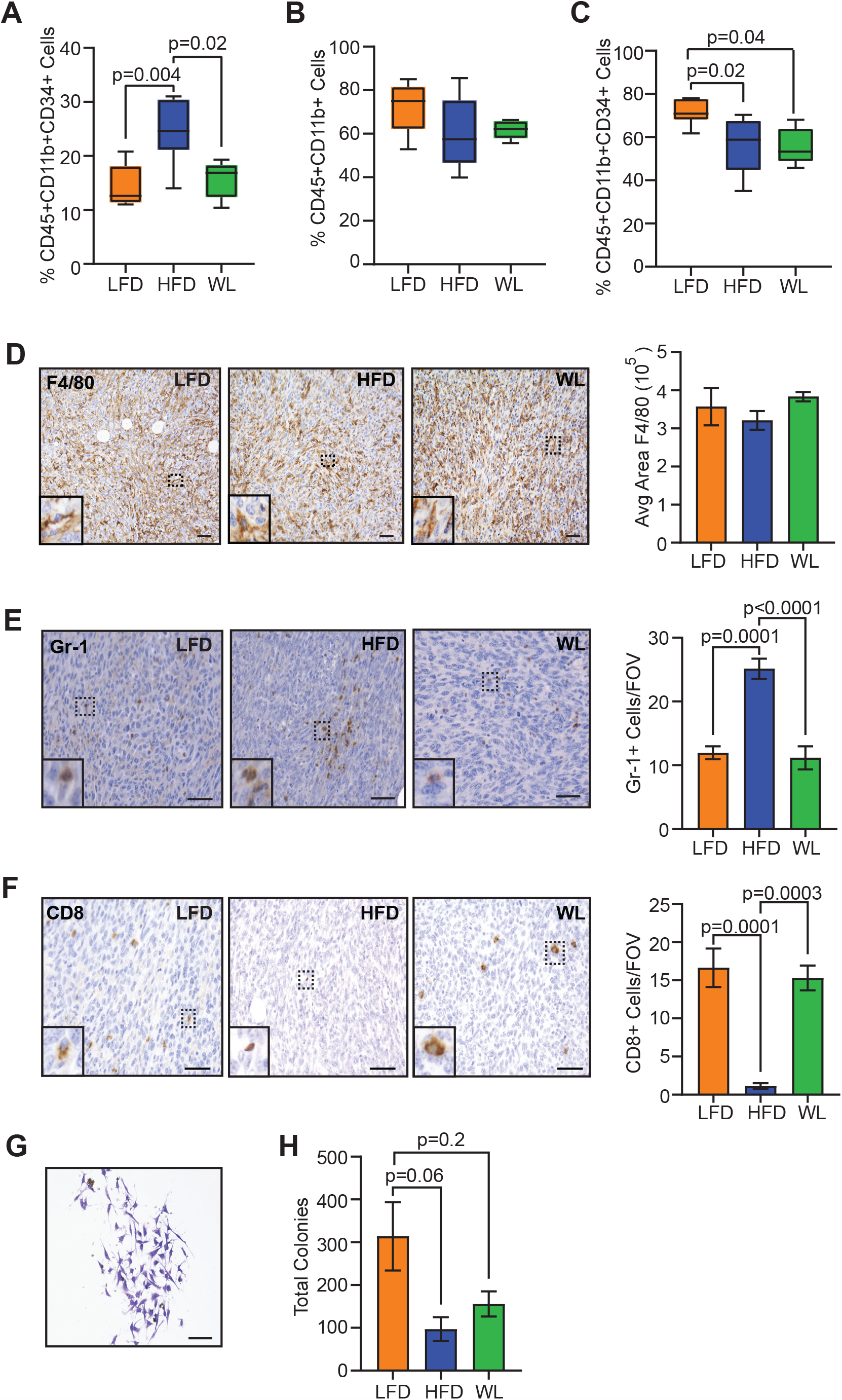
Weight loss reduces immunosuppressive microenvironment within tumors. **(A)** CD45^+^CD11b^+^CD34^+^ myeloid progenitor cells in bone marrow quantified by flow cytometry (n = 6/group). **(B)** CD45^+^CD11b^+^ myeloid cells in TC2 tumors quantified by flow cytometry (n = 6/group). **(C)** CD45^+^CD11b^+^CD34^+^ immature myeloid cells in TC2 tumors mice quantified by flow cytometry (n = 6/group). **(D)** Representative images and quantification of F4/80^+^ cells in TC2 tumors (n=6/group). **(E)** Representative images and quantification of Gr-1^+^ cells in TC2 tumors (n=6/group). **(F)** Representative images and quantification of CD8^+^ cells in TC2 tumors (n=6/group). **(G)** Representative crystal violet image of fibrocyte colony from CD11b^+^ cells isolated from TC2 tumors. **(H)** Quantification of total fibrocytes colonies from CD11b^+^ cells isolated from TC2 tumors (n=4/group). Magnification bar = 50 μm.

To assess myeloid lineage cells within tumors, we stained tumor sections from lean, obese, and formerly obese mice with antibodies to detect macrophages and myeloid-derived suppressor cells (MDSC). Although F4/80^+^ macrophages made up a large component of the tumor microenvironment, no significant differences were detected among the tumors from mice in each group (Figure 4D). However, Gr1^+^ MDSC were significantly increased in the tumors of obese mice compared to lean mice (p=0.0001, Figure 4E), and formerly obese mice had significantly reduced MDSC in tumors compared to obese mice (p<0.0001, Figure 4E). MDSC have been characterized as immunosuppressive within tumors [37, 38]. Consistent with this immunosuppression, CD8^+^ T cells were significantly reduced in tumors from obese mice compared to tumors from either lean mice (p=0.0001) or formerly obese mice (p=0.0003, Figure 4F). These results suggest that obesity creates an immunosuppressive tumor microenvironment, while weight loss prior to tumor growth reverses these effects.

Since we observed increased collagen deposition and CAF in tumors from obese mice, we hypothesized that obesity may also increase fibrocytes in the myeloid cell populations. To test this hypothesis, we isolated CD11b^+^ cells from the tumors of lean, obese, and formerly obese mice and quantified colony formation *in vitro* (Figure 4G). Interestingly, we observed significantly less total colony formation from the CD11b^+^ cells isolated from tumors (Figure 4H) than from mammary glands (Figure 2F). In contrast to our hypothesis, we observed reduced fibrocyte colonies in CD11b^+^ cells isolated from tumors of obese mice compared to lean mice (p=0.06, Figure 4H), while fibrocyte colonies formation were not significantly different among tumors of lean and formerly obese mice (p=0.2, Figure 4H). These results suggest that differentiation of immature myeloid cells into fibrocytes is reduced in the tumor microenvironment.

### Myeloid progenitor cells contribute to collagen deposition in mammary tumors of lean and obese mice

We have previously observed that immature myeloid cells isolated from the mammary glands of obese mice expressed collagen 1 and collagen 3 [21], which are believed to be defining features of fibrocytes that are absent from other hematopoietic cells [39]. Since we observed increased collagen deposition and CAF within the tumors of obese mice, we hypothesized that fibrocytes present within the mammary glands during tumor formation may promote a more fibrotic tumor microenvironment. To test this hypothesis, EGFP^+^ FVB/N mice, which ubiquitiously express enhanced GFP [27], were fed HFD for 16 weeks to induce obesity. Using FACS, we isolated live CD45^+^ cells, CD11b^+^CD34^-^ monocytes, and CD11b^+^CD34^+^ myeloid progenitor cells, which are the cells of origin of fibrocytes, from bone marrow of obese EGFP^+^ donor mice. TC2 mammary tumor cells were mixed with the sorted bone marrow populations, then injected into the mammary glands of lean and obese mice (Figure 5A). When transplanted with tumor cells, the bone marrow populations did not alter tumor growth in either lean or obese mice (Figure 5B, C).

**Figure 5.**
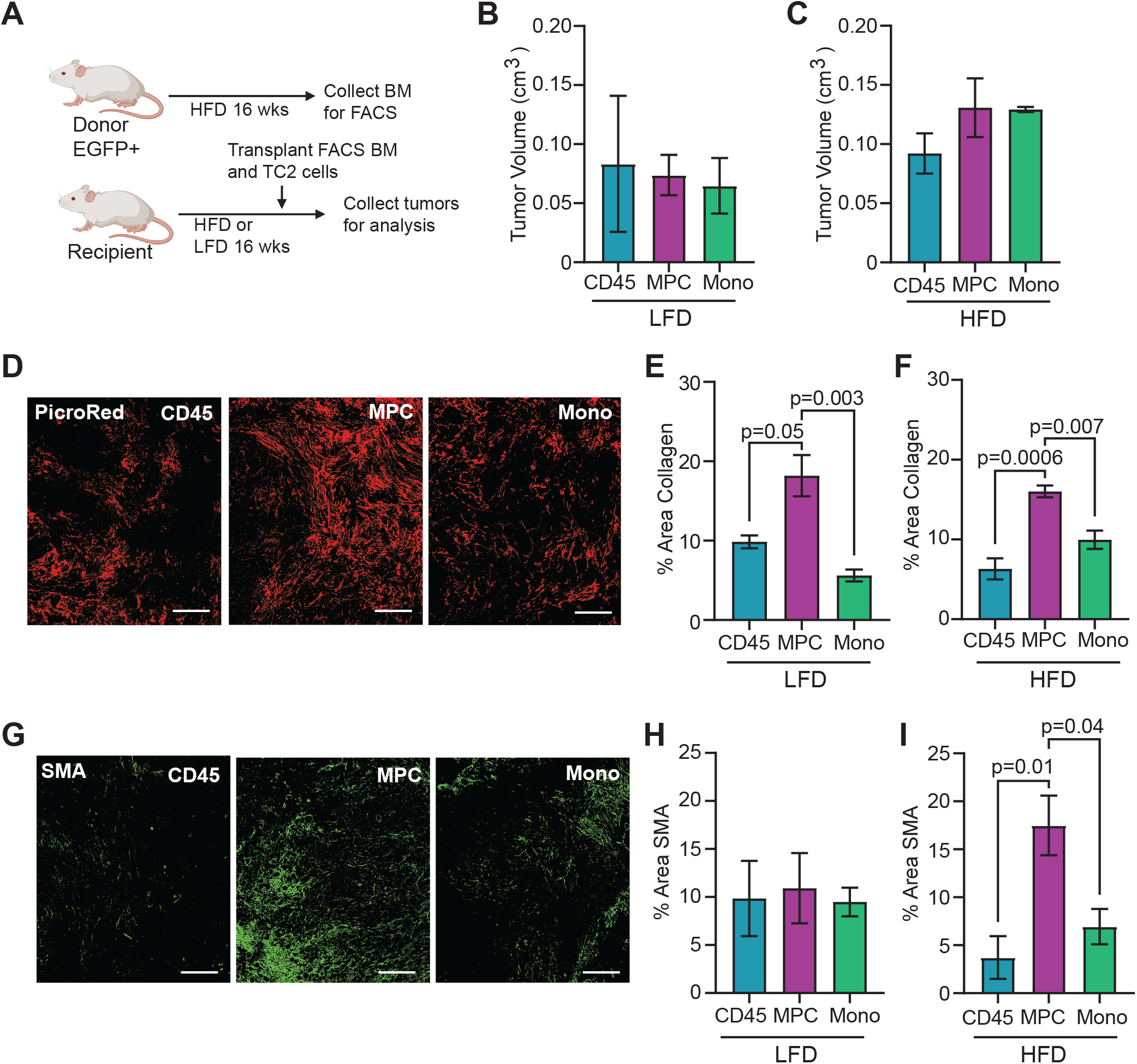
Myeloid progenitor cells contribute to TC2 mammary tumor collagen deposition, but not tumor growth. **(A)** Schematic of tumor transplant experiment. **(B)** Tumor volume of lean mice on Day 36 (n = 4/group; Kruskal-Wallis with Dunn’s multiple comparisons test). **(C)** Tumor volume of HFD-fed mice on Day 25 (n = 4/group; Kruskal-Wallis with Dunn’s multiple comparisons test). **(D)** Representative images of picrosirius red-stained collagen in mammary tumors isolated from obese mice. **(E)** Collagen within tumors normalized to total tumor area from lean mice (n=4/group). **(F)** Collagen within tumors normalized to total tumor area from obese mice (n=4/group). **(G)** Representative images of SMA expression in mammary tumors from obese mice. **(H)** SMA expression within tumors normalized to total tumor area from lean mice (n=4/group). **(I)** SMA expression within tumors normalized to total tumor area from obese mice (n=4/group). Magnification bar: **(D, G)** 300 μm.

To assess the impact of the isolated bone marrow cell populations on the mammary tumor microenvironment, collagen deposition was quantified within tumors from each group using picrosirius red staining (Figure 5D). Transplant of myeloid progenitor cells with the tumor cells significantly increased collagen within tumors from both lean (p=0.003, Figure 5E) and obese (p=0.007, Figure 5F) mice, compared to the more differentiated CD11b^+^CD34^-^ population of monocytes. To quantify CAF, tumors from all groups were stained for SMA (Figure 5G). While no significant differences were observed in tumors from lean mice (Figure 5H), SMA was significantly increased in tumors grown with myeloid progenitor cells from obese mice, compared to those transplanted with CD45^+^ cells (p=0.01) or CD11b^+^CD34^-^ monocytes (p=0.04, Figure 5I). Together, these results suggest that fibrocytes present at the time of tumor formation promote the development of desmoplastic stroma in obese mice.

Fibrocytes have been shown to differentiate into myofibroblasts in models of other types of cancer [40, 41] or stimulate other cells to promote fibrosis [42, 43]. To identify whether the transplanted EGFP^+^ cells contributed to the CAF population, we dissociated tumors and quantified GFP expression using flow cytometry. We did not detect a significant number of GFP^+^ cells in the tumors from any group of mice (Figure S3). These data suggest that the transplanted bone marrow cells did not expand in number to significantly contribute to the CAF population but may instead promote the differentiation or recruitment of other stromal cells to become CAF.

## DISCUSSION

Obesity is associated with poor breast cancer prognosis [6]. While weight loss improves outcomes for other health conditions, little was known about how weight loss prior to tumor formation potentially impacted the growth and microenvironment of mammary tumors. Our studies suggest that while weight loss did not completely reduce the rate of tumor growth, the microenvironment of the resulting tumors was less fibrotic and immunosuppressive than those from obese mice. While fibrocytes appear to be less frequent in the growing tumors, their presence in the mammary gland prior to tumor formation may promote the rapid formation of CAF in the early tumor microenvironment. We observed that co-mixing of myeloid progenitor cells from obese mice with TC2 tumors cells resulted in significantly increased collagen deposition in tumors of both lean and obese mice. Fibrocytes may promote fibrotic changes in resident fibroblasts and adipose-derived stromal cells to become CAF in the developing tumor microenvironment, leading to more desmoplastic tumors observed clinically in breast cancer patients with obesity [8].

Within the mammary glands of non-tumor-bearing mice, weight loss did not resolve the increased collagen deposition around mammary ducts, indicating that fibrosis may be a longer-lasting microenvironmental condition than inflammation. These results are consistent with human studies of subcutaneous and visceral fat following weight loss through bariatric surgery [15, 16]. The presence of elevated collagen in the mammary glands of formerly obese mice may reflect slower tissue remodeling of mature collagen fibers within the tissue [44]. Consistent with a decrease in myeloid progenitor cells in the bone marrow, we observed a decrease in fibrocytes within the mammary glands of formerly obese mice. These results suggest that with the resolution of the chronic inflammation of obesity, fibrocytes are not continually recruited to the tissue.

While we observed that weight loss reduced CLS, which are a histological marker for local inflammation [45], the total CD11b^+^ cell population was not reduced. Studies of macrophages within different adipose tissue depots have demonstrated that the macrophage population is heterogeneous, depending on microenvironment conditions [46-48]. Macrophages that form CLS may be functionally distinct from macrophages in other locations within adipose tissue [49]. Macrophages can also acquire a metabolically activated phenotype [50] due to removal of lipid from dying adipocytes [51]. In visceral fat during weight loss, macrophage populations shift to include those with a phagocytotic function, which may participate in tissue remodeling [47, 52]. CD11b is expressed at various levels by multiple different cell types [53], and additional gene expression experiments and expanded flow cytometry markers are required to characterize the distinct populations of CD11b^+^ cells present in the mammary glands of lean, obese, and formerly obese mice.

Similar to the CD11b^+^ cell population in the mammary gland, the CD11b^+^ cell population in mammary tumors is heterogeneous. The growing mammary tumor may affect both the types and functions of myeloid cells within the tumor microenvironment [54]. Further, mammary tumors can select for populations of myeloid cells within bone marrow [55, 56], which changes the composition of myeloid cells recruited into tumors. Although we observed an increase in fibrocytes within the immature myeloid cell population in the mammary glands of obese non-tumor-bearing mice, fibrocytes were no longer enhanced within this population in the tumor microenvironment. Cytokines and growth factors secreted by tumor cells may shift the differentiation of immature myeloid cells into MDSC and tumor-associated macrophages [57]. Consistent with this idea, we observed increased Gr-1^+^ MDSC in the tumors of obese mice, and the immunosuppressive environment was reflected in significantly reduced CD8^+^ T cells. GM-CSF is one cytokine that has been implicated promoting the differentiation of MDSC from immature myeloid cells within tumors [58, 59]. Obesity may enhance the effects of the tumor cells by expanding the myeloid progenitor cell population prior to tumor formation.

Increasing desmoplasia, including SMA^+^ stromal cells, are associated with worse survival in breast cancer patients [60, 61]. Weight loss is a commonly recommended intervention for obesity and may reduce breast cancer risk [10, 11]. Here we show that weight loss prior to ERα^+^ tumor formation limits the desmoplasia and immunosuppression within mammary tumors, which may suggest improved breast cancer outcomes. In contrast, in models of triple negative breast cancer, mice that lost weight due to decreased consumption of a HFD developed tumors with aggressive characteristics that were more similar to those from obese mice [62, 63]. It is possible that changes in the mammary microenvironment due to weight loss may have divergent effects on different subtypes of breast cancer. Further, alternate methods of weight loss may impact the resulting mammary tumors in different ways. Following injection with EO771 tumor cells, mice that lost weight through low-fat calorie restriction, Mediterranean-style calorie restriction, and intermittent-calorie restriction had reduced tumor growth and least expression of genes associated with epithelial-to-mesenchymal transition [62]. In addition, in a model of weight loss through bariatric surgery, tumors that developed in the surgical group had higher expression of genes associated with an inflammatory response and improved responses to anti-PD-L1 immune checkpoint therapy [64]. Together these studies suggest that the method of weight loss may have long-term impact on the biology of the tumors that develop following weight loss. Understanding how different methods of weight loss alter the microenvironment and biology of tumors may lead to improved prevention strategies for women at high risk for breast cancer.

## Acknowledgements

The authors would like to thank Brenna Walton and Mason McGuire for critically reading the manuscript and helpful discussions. This research was supported by NIH F31CA247265 (GK) and R01CA227542 (LMA). The authors would like to acknowledge the P30CA014520-UW Carbone Cancer Center Support Grant (CCSG).

## Authors’ contributions

G. Kuziel, B.N. Moore, G.P. Haugstad, Y. Xiong, and A.E. Williams collected and analyzed the data, and interpreted the results. G. Kuziel drafted the manuscript. G. Kuziel and L.M. Arendt conceived of the study, designed the experiments, interpreted the results, wrote the manuscript, and provided financial support. All authors read and approved the final manuscript.

