## Supplementary Table 1 and Supplementary Figures 1-3 for "Alterations in the Mammary Gland and Tumor Microenvironment of Formerly Obese Mice"

**Supplementary Tables**

**Table S1.** Antibodies used for flow cytometry, immunofluorescence, and immunohistochemistry.

| | Concentration | Catalog Number | Supplier | RRID |
|---|---|---|---|---|
| Fixable Viability Dye eFluor780 | 1:1000 | 65-0865 | ThermoFisher | |
| CD16/32 | 0.5 ng/µl | 14-0161-85 | ThermoFisher | AB_467133 |
| BV421 CD34 | 16 ng/µl | 562608 | BD Biosciences | AB_11154576 |
| APC CD11b | 2.5 ng/µl | 17-0112-82 | ThermoFisher | AB_469343 |
| PE CD45 | 2.5 ng/µl | 12-0451-82 | ThermoFisher | AB_465668 |
| CD11b | 0.1 µg/µl | 14-0112-85 | ThermoFisher | AB_467108 |
| F4/80 | 1:250 | 123102 | BioLegend | AB_893506 |
| CD11b | 1:200 | NB110-89474 | Novus | AB_1216361 |
| α-Smooth Muscle Actin (SMA) | 1:1500 | A5228 | Sigma-Aldrich | AB_262054 |
| Collagen I | 1:200 | NB600-408 | Novus | AB_10000511 |
| Estrogen Receptor α | 1:500 | 06-935 | EMD Millipore | AB_310305 |
| Ly-6G / Ly-6C | 1:200 | MA1-10401 | ThermoFisher | AB_11152791 |
| CD8 | 1:200 | NBP1-49045 | Novus | AB_10011327 |
| 488 anti-Mouse | 1:300 | A11001 | ThermoFisher | AB_2534069 |
| 488 anti-Goat | 1:300 | A11055 | ThermoFisher | AB_2534102 |
| 546 anti-Mouse | 1:300 | A11003 | ThermoFisher | AB_2534071 |
| 546 anti-Rabbit | 1:300 | A11010 | ThermoFisher | AB_2534077 |
| Biotinylated anti-Rabbit | 1:500 | BA-1000 | Vector | AB_2313606 |
| Biotinylated anti-Rat | 1:250 | BA-4000 | Vector | AB_2336206 |

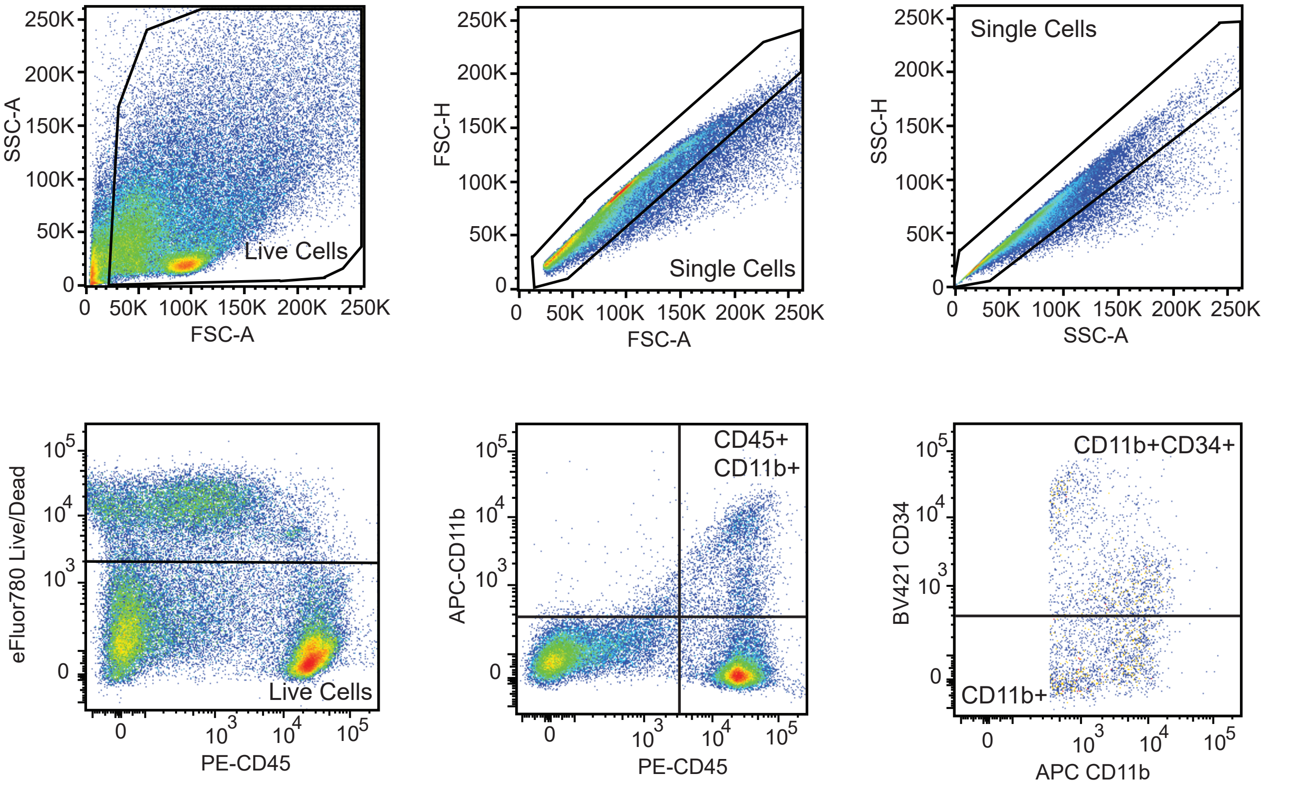


**Supplementary Figure 1.** Flow cytometry gating strategy of mammary gland. Cells were gated for debris, followed by single cells, and live cells using viability dye. Live cells were gated for CD45 and CD11b expression, and CD11b+ cells were further gated for CD34 expression.

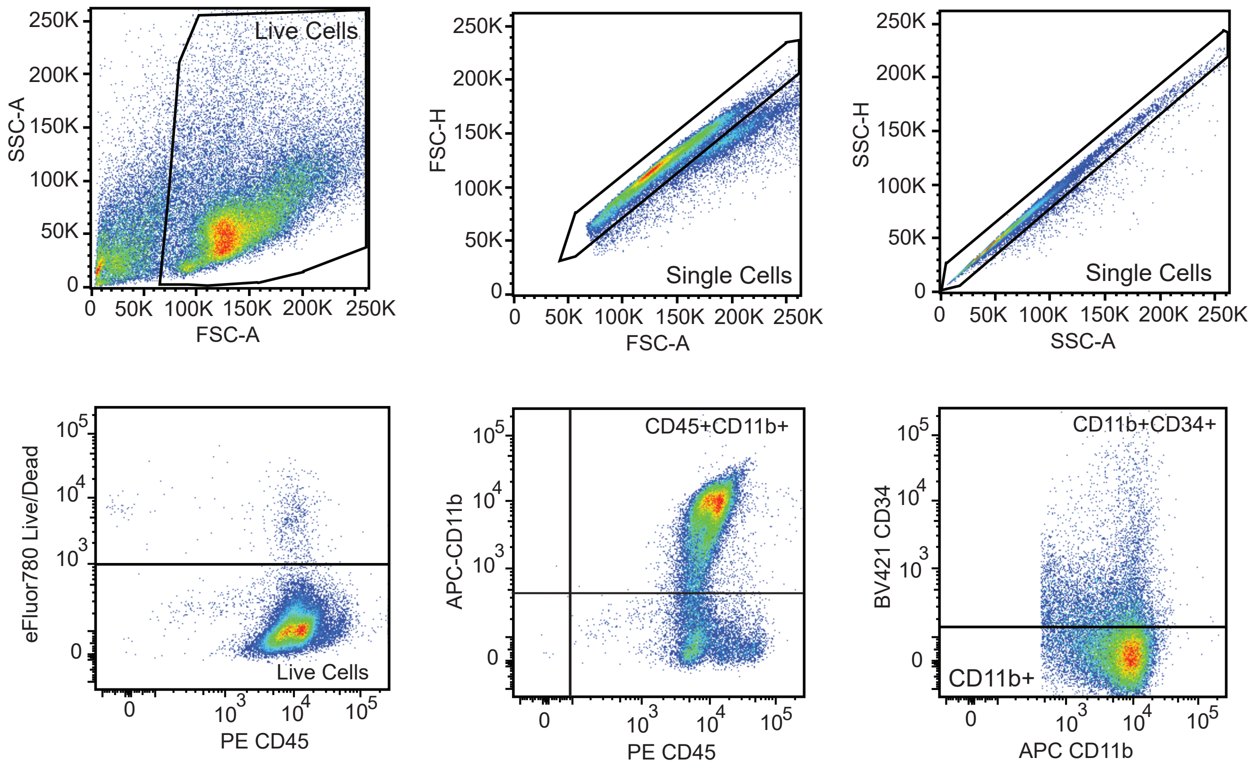


**Supplementary Figure 2.** Flow cytometry gating strategy of bone marrow. Cells were gated for debris, followed by single cells, and live cells using viability dye. Live cells were gated for CD45 and CD11b expression, and CD11b+ cells were further gated for CD34 expression.

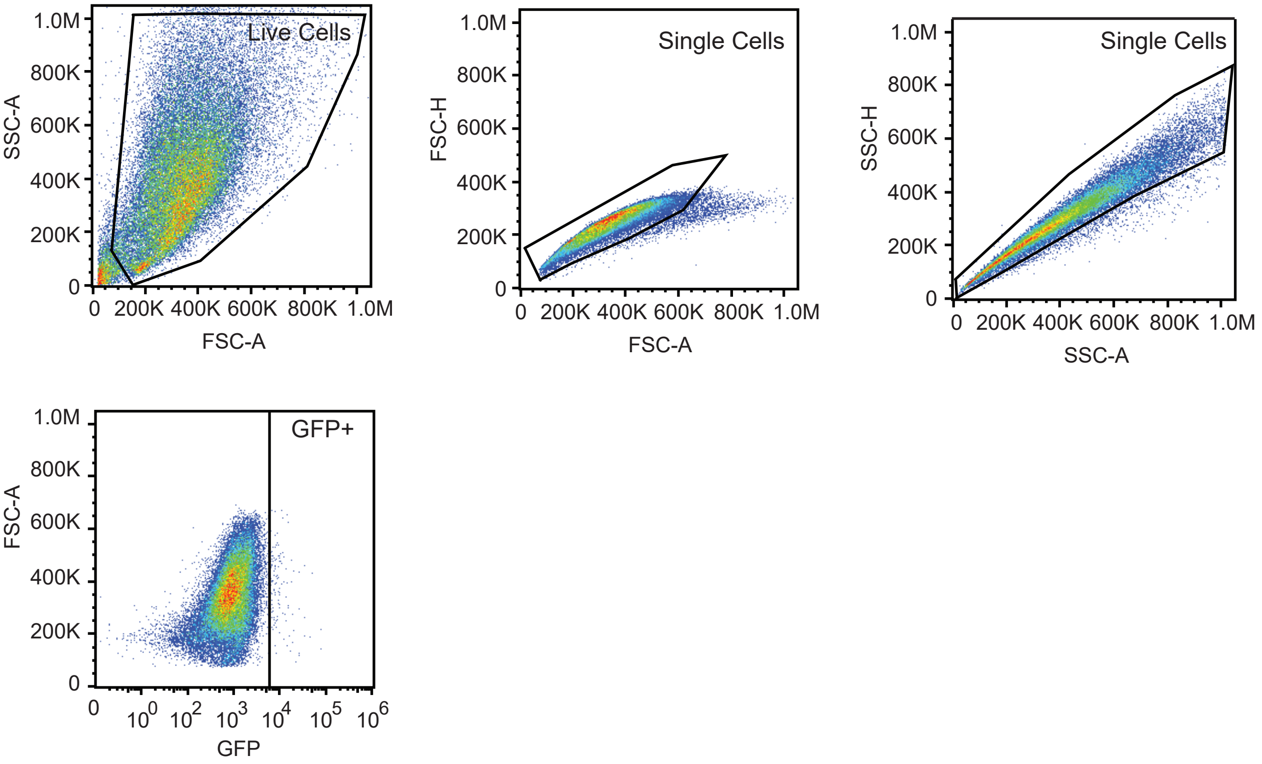


**Supplementary Figure 3.** Flow cytometry gating strategy to detect GFP in TC2 tumors mixed with bone marrow cell populations. Cells were gated for debris, followed by single cells, and then gated for GFP expression.
